# A proteogenomic approach to discover novel lncRNA-derived peptides and their potential clinical utility in hepatocellular carcinoma

**DOI:** 10.1101/2025.09.30.679478

**Authors:** Bingwu Li, Kandarp Joshi, Dan Ohtan Wang

## Abstract

Peptides are increasingly recognized for their versatile functions in biological contexts but their clinical relevance and utility remain largely unexplored. Proteogenomic approaches can accelerate peptide discovery in clinical samples by integrating proteomic data with genomics and transcriptomics evidence. However, long noncoding RNA (lncRNA)-derived peptides (lncPeps) remain largely unidentified, resulting in unmatchable MS/MS spectra. To solve this problem, we have used high-quality Ribo-seq translatomic datasets to generate an extensive database of human liver lncPeps, which we subsequently applied to proteomics data of tumor–adjacent normal tissue pairs from hepatocellular carcinoma (HCC) patients. Using the new database, we discovered 105 novel lncPeps including lncPeps differentially expressed between tumor and non-tumor tissues, and lncPeps with significant correlation with prognosis. Remarkably, combining the expression of lncPeps with canonical proteins in a LASSO regression model improved predictive performance for recurrence, increasing the AUC by 0.005 to 0.085 across three recurrence time points. These findings suggest that lncPeps discovery contributes to our understanding of the molecular heterogeneity and progression of HCC, and broadens the range of potential biomarker candidates or treatment targets for the disease.

## Introduction

Hepatocellular carcinoma (HCC) is the third leading cause of cancer-related mortality worldwide and the fastest-rising cause of cancer-associated deaths in Western countries (1, 2). Although surgical treatment can be effective in the early stages, once HCC develops, the overall five-year survival rate remains at only 50–70% (1). With advances in multi-omics and gene-editing technologies, precision medicine has brought new hopes to patients with HCC. However, the high degree of molecular heterogeneity in HCC presents numerous challenges, including low response rates, drug resistance, and immune evasion (3). To overcome these issues, it is imperative to gain a thorough understanding of the molecular mechanisms underlying tumor heterogeneity. Furthermore, early diagnosis of HCC remains problematic, and the development of clinically useful biomarkers for HCC management has been slow. Alpha-fetoprotein (AFP) has been used as a biomarker in clinical practice for over six decades; yet its use in diagnosis remains controversial due to variable sensitivity and specificity (4, 5).

Proteomic analyses have provided critical insights into biomarker discovery and therapeutic strategies. A deeper understanding of the human proteome is essential for deciphering the molecular mechanisms of cancers such as HCC and for advancing precision medicine (6–8). Despite extensive efforts over the years, a substantial portion of the proteome remains uncharacterized (9). Expanding systematic annotation and functional characterization are crucial for identifying previously unrecognized protein-coding elements, particularly short peptide sequences (with sequence length less than 100 aa) that had been traditionally classified as “noncoding”. A large fraction of these unannotated peptides may be nested within lncRNAs. In SmProt, a depository of small proteins with experimental evidence of translation, 10.2% of the small ORFs (sORFs) are nested in lncRNA. The number of peptides translated from lncRNAs is likely underestimated due to insufficient annotation of lncRNA and ORFs (10), despite the fact that lncPeps show physiological significance in numerous biological contexts including cancer. Several lncPeps have been shown to modulate tumor growth and metastasis by interacting with oncogenic signaling pathways or tumor suppressors (11–15). In addition, lncPeps have been implicated in immune regulation (16, 17) and contribute to chemotherapy resistance by altering the tumor microenvironment and affecting drug response (17–21).

Recent reports have also demonstrated the potential of lncPeps as biomarkers and therapeutic targets for HCC, offering unique insights into HCC pathogenesis and molecularly targeted therapy. Guo *et al*. computationally predicted an sORF in lncRNA *ZFAS1* using machine learning, which encodes a micropeptide that elevates intracellular reactive oxygen species (ROS) production by inhibiting nicotinamide adenine dinucleotide (NAD) dehydrogenase expression and thus promotes cancer cell migration (22). Micropeptide in mitochondria (MPM) interacts with NAD dehydrogenase NDUFA7, inhibiting Mitochondrial complex I activity and ultimately inhibiting the metastasis of HCC cells (23). lncPep KRASIM was identified using Ribo-seq, which interacts with the oncogenic protein KRAS and suppresses HCC cell growth and proliferation (24). lncPep PINT87aa was identified by Ribosome Nascent-chain Complex RNA sequencing (RNC-seq). PINT87aa directly binds to FOXM1 to block PHB2 transcription, inducing cell cycle arrest and cellular senescence thus playing a tumor-suppressive role (25). lncPep SMIM30 was identified by RIP-seq assay with an antibody against ribosomal protein S6 (RPS6). SMIM30 functions as an adaptor for the membrane anchoring of SRC/YES1 and contributes to its activation and promotes HCC tumorigenesis (26). lncPep Linc013026-68AA, identified through qRT-PCR analysis of polysome fractions, is suggested to play a role in regulating cell proliferation, although its precise mechanism of action remains unclear (27). Another lncRNA, *AC115619*, encodes a peptide of 22 amino acids (AC115619-22aa) that has been proposed to disrupt the formation of N6-methyladenosine (m6A) methylation complexes, leading to reduced global m6A levels and HCC progression (28). Additionally, the mitochondrial RNase P inhibitory peptide (MRPIP), derived from an lncRNA *AC027045.3*, has been shown to suppress HCC progression by modulating mitochondrial RNA processing pathways (29).

However, lncPeps in the human genome remain largely unannotated by conventional genome annotation tools owing to their noncanonical and noisy nature in proteomic data (30). To resolve this problem, proteogenomic analyses are increasingly employed for aiding noncanonical peptide discovery by integrating genomic, transcriptomic, and proteomic data (31), but remain controversial due to high false positive rates (32). More recently, the incorporation of translatomics data—particularly Ribo-seq—has been emphasized as an effective strategy to reduce false positives (33, 34). Ribo-seq, a representative translatomics technique, has played a crucial role in noncanonical peptide discovery (35).

Motivated by the recent expansion in lncRNA annotation (NONCODE V6) and more advanced proteomic techniques (6, 36), we explored hepatocellular carcinoma (HCC)-relevant lncPeps with a new proteogenomic pipeline that integrates Ribo-seq and proteomic datasets. We initially collected high-quality human Ribo-seq datasets including 5 cancerous and 2 paracancerous liver tissues, as well as 5 biological samples from primary human hepatocytes and 5 samples from two liver cancer cell lines. To identify lncRNA-ORFs, we used GENCODE (version 46) and NONCODE (version 6) as references and three ORF identifiers to generate lncRNA-ORFs indices (37, 38). The identified lncRNA-ORFs were translated into a lncPep database for lncPep discovery in HCC patients. Biopsy proteomic data from cancerous/paracancerous paired tissues of 268 patients, from two published cohorts, were retrieved from published databases and analyzed. (6, 36). We identify lncPeps differentially expressed in cancerous tissues (e.g., *HNRNPA1P36*), with a subset (including *PPIAP79* and *POTEKP*) also exhibiting significant correlations with patient survival and recurrence prognosis. This study demonstrates the potential of using proteogenomic pipeline to discover functional lncPeps in HCC and possibly other cancers.

## Experimental procedures

### Ribo-seq data collection and alignment, and quality control

Ribo-seq datasets were downloaded from the NCBI GEO database (Supplementary Table 1) (39). Trim Galore was used for preprocessing to remove adapter sequences and low-quality reads (40). Human rRNA sequences were downloaded from the Rfam database, and bowtie2 was used to remove rRNA reads (41, 42). The remaining reads were aligned against the human reference genome hg38 using STAR with the following parameters: ‘--outFilterType BySJout --outFilterMismatchNmax 2 --outSAMtype BAM SortedByCoordinate --quantMode TranscriptomeSAM –outFilterMultimapNmax 1 --outFilterMatchNmin 16 --readFilesCommand zcat --outReadsUnmapped None --alignEndsType EndToEnd’ (43). Multi-mappers were discarded.

All Ribo-seq data underwent rigorous quality control, with the methods and criteria detailed below. The metaplot function from RiboCode and the quality function from Ribo-TISH were employed to analyze the ribosome-protected fragments (RPFs) (44, 45). The criteria for quality control required RPF reads to be 27–30 nucleotides in length, exhibit triplet periodicity within the coding sequence (CDS) region as determined by both RiboCode and Ribo-TISH and have a P-site offset of 12 nucleotides. Only reads meeting these strict criteria were used for open reading frame (ORF) prediction and quantification. After applying these filters, 17 datasets were selected for further analysis (Supplementary Table 1).

BEDTools intersect function was used to analyze the distribution of RPF reads across different RNA features (46). Reads mapped to genomic features were counted using the *featureCount* function from the Subread package and normalized to FPKM value (47, 48).

### Assessment of translational potentials of open reading frames on lncRNAs

lncRNA annotation files were obtained from GENCODE (Version 46) and NONCODE (Version 6) as reference files for annotating human lncRNA-ORFs (37, 38). RiboCode, Ribo-TISH, and ribotricer were used to detect actively translating ORFs (44, 45, 49). Across all three tools, the minimum ORF length was set to 18 nucleotides. For translation initiation, ATG, CTG, GTG, and TTG were considered as potential start codons, while TAG, TAA, and TGA were used as stop codons for translational termination. For RiboCode and Ribo-TISH, both the longest strategy and frame-best strategy were used to predict ORFs, with the default P-value (Stouffer’s method combined P-value < 0.05) employed for identifying actively translating ORFs. In the case of ribotricer, the recommended human phase-score cutoff was applied for active ORF prediction, along with a minimum ratio of codons with non-zero reads set to 0.4.

The coding potential of transcripts was calculated by CPC2 (standalone version) (50), CPC2 assigns a score ranging from 0 to 1, with higher values indicating greater similarity between the input transcript and known mRNA sequences, and a higher likelihood of protein-coding capacity. The RPF track for a specific ORF is plotted by RiboCode’s *plot_orf_density* function.

### Building a custom proteogenomics database

The active ORFs predicted by the three tools were merged and collapsed, with ORFs sharing the same stop codon across different transcript isoforms being retained as the longest ORF or the best ORF (based on the minimum P-value).

Based on the Ribo-seq results, we constructed three proteogenomic database candidates with varying levels of stringency. Database 1 included lncPeps predicted by two or more of the three tools, totaling 9,451 lncPeps (RiboCode ∩ Ribo-TISH) ∪ (RiboCode ∩ ribotricer) ∪ (Ribo-TISH ∩ ribotricer). Database 2 included lncPeps predicted by at least one tool, totaling 33,083 lncPeps (RiboCode ∪ Ribo-TISH ∪ ribotricer). Database 3 included all candidate ORFs from transcripts showing RPF signals. In this case, no triplet periodicity filtering was applied, resulting in 288,383 ORFs. These databases reflect different levels of tolerance in defining lncRNA-ORF pools. Additionally, lncRNA coding peptides annotated by GENCODE (V46) and UniProt (Swiss-Prot) were added (38, 51). To facilitate group FDR estimation, we customized the FASTA headers for the lncRNA coding peptide database to distinguish them from canonical proteins.

### Computational proteomics analysis

Proteomic data from hepatocellular carcinoma patient biopsies were obtained from the IProX and CPTAC repositories, as reported by Jiang *et al*. and Gao *et al*. (6, 36, 52). Using the custom lncPep indices along with the UniProt (Swiss-Prot) human protein database, we annotated the datasets following analysis with the FragPipe computational platform (version 22). MSFragger was used for peptide identification, trypsin was selected as the digestion enzyme, allowing up to two missed cleavage sites. Cysteine carbamidomethylation was set as a fixed modification, while N-terminal acetylation and methionine oxidation were considered variable modifications. The mass tolerance was set to 20 ppm for the initial search and 4.5 ppm for the main search. Known laboratory contaminants were included in the analysis. A reverse-sequence decoy database was generated and used for false discovery rate (FDR) estimation (53).

DIA-NN was utilized to predict MS2 spectra, retention time (RT), and ion mobility (IM), while MSBooster integrated peptide-spectra matching features with deep learning-based predictions. Philosopher was used for false discovery rate (FDR) estimation (54, 55). To evaluate FDR estimation for lncRNA-encoded peptides, we used two strategies: group-specific FDR estimation and the two-pass search (56, 57). In the group-specific FDR estimation, lncRNA-encoded peptides and known proteins were analyzed together, but the FDR was calculated separately for each group. In the two-pass search strategy, known proteins were first subjected to FDR estimation. Spectra that did not pass the FDR threshold was then used to search for lncRNA-encoded peptides, after which the spectra containing these peptides were subjected to a separate FDR estimation. In both methods, the FDR threshold was set to 1% (58).

All lncRNA-derived peptide fragments were mapped against previously annotated proteins in UniProtKB and GENCODE. Peptide fragments that fully mapped to known proteins were excluded. Peptide fragments with a single amino acid mismatch were also removed as they were considered potential single amino acid polymorphism (SAP) peptides.

The PDV tool was used as visualization of the mass spectrometry (59).

### Quantification of the HCC-lncPeps

The resulting high-confidence lncPeps spectra were used for quantitative analysis, with the MSstats and MSstatsTMT packages applied for protein abundance estimation and normalization of protein quantification results for label-free (LBF) and tandem mass tag (TMT) data, respectively (60, 61). For LBF data, Tukey’s median polish was used for protein abundance estimation, and the equalizeMedians method was applied for normalization. In the case of TMT data, the MSstats method was used for protein abundance estimation, with normalization based on the reference channel. The *removeBatchEffect* function from the limma package was utilized for batch effect removal in TMT quantification results, and principal component analysis (PCA) plots were generated to assess batch effects (62). In all approaches, lncRNA-encoded peptides were analyzed alongside canonical proteins.

To address missing values in the expression matrix, we applied two strategies: filtering out samples with missing values and applying minimal imputation, with both approaches included in subsequent analyses.

### Biostatistical analysis

Logistic regression analysis was performed using *glm* function in the stats package. Cox regression analysis for survival analysis was performed using *coxph* function and *survdiff* function was used for log-rank test from survival package (63, 64). For the log-rank test, patients were stratified by lncPep expression into quartiles. The upper quartile (top 25%) was defined as the high-expression group, and the lower quartile (bottom 25%) as the low-expression group.

For RNA-level analyses, all TCGA data were collected from the UCSC Toil RNA-seq Recompute project (65), TPM values were used as normalized gene expression, and missing values were imputed using zero.

### Functional predictions

GSEA analysis was performed using the *gseGO* function from the clusterProfiler package. For the lncPeps of interest, Pearson correlation coefficients were calculated between their intensities and those of all known proteins. The known proteins were then ranked based on these Pearson correlation coefficients, and GSEA was conducted using *gseGO* with the Gene Ontology (GO) database (66). GO terms with an adjusted p-value less than 0.05, corrected using the Benjamini-Hochberg method, were considered significantly enriched.

### Machine Learning Modeling

For the MS-peptide (lncPeps derived from Ribo-seq data that have been detected by proteomic mass spectrometry) vs. Non-MS-peptide (lncPeps derived from Ribo-seq data but not detected by proteomic mass spectrometry) classification model, the seqinr package and the AAindex database were utilized to extract the physicochemical properties of lncPep (67, 68). The caret package was employed for data partitioning and machine learning, with 70% of the data randomly sampled as the training set. A LASSO regression model was used for feature selection: the glmnet package implemented LASSO, the *cv.glmnet* function performed 10-fold cross-validation, and the glmnet function fit a generalized linear model via penalized maximum likelihood. The regularization parameter was determined based on lambda.min (69). The final machine learning model was built using the Random Forest algorithm (70). To address the imbalance in the numbers of MS-peptides and Non-MS-peptide during model training, the *downSample* function from the caret package was used for downsampling, and 10 rounds of downsampling were performed to mitigate random sampling bias (70). Additionally, the *Boruta* function in the Boruta package was used to calculate the variable importance measure (VIM) for the LASSO-filtered features in each downsampling (71). The pROC package was used for plotting the ROC curves (72).

For the cancer tissue classification and prognosis prediction models, the normalized expression of canonical proteins and lncPeps was used as input features. The same feature selection method described above was employed. To enhance the interpretability of the model, we utilized a simpler LASSO regression for modeling, without performing any sub-sampling during training. To compare the model based solely on canonical protein expression with the one incorporating both canonical protein and lncPep expression, a likelihood ratio test was conducted using the *anova* function in the stats package of R.

## Results

### Systematic assessment of lncRNA-ORFs in human liver using coding potentials

The proteogenomic strategy employed in this study is illustrated in Figure 1a. In brief, the workflow comprises two components: translatomics and proteomics analysis. To systematically investigate lncRNA-derived ORFs (lncRNA-ORFs) in human liver tissues and cells, we constructed a liver lncRNA-ORFs database based on quality-controlled Ribo-seq data derived from relevant biological samples. This database was subsequently used to guide database searches and peptide identification from MS/MS proteomic datasets.

**Figure 1:**
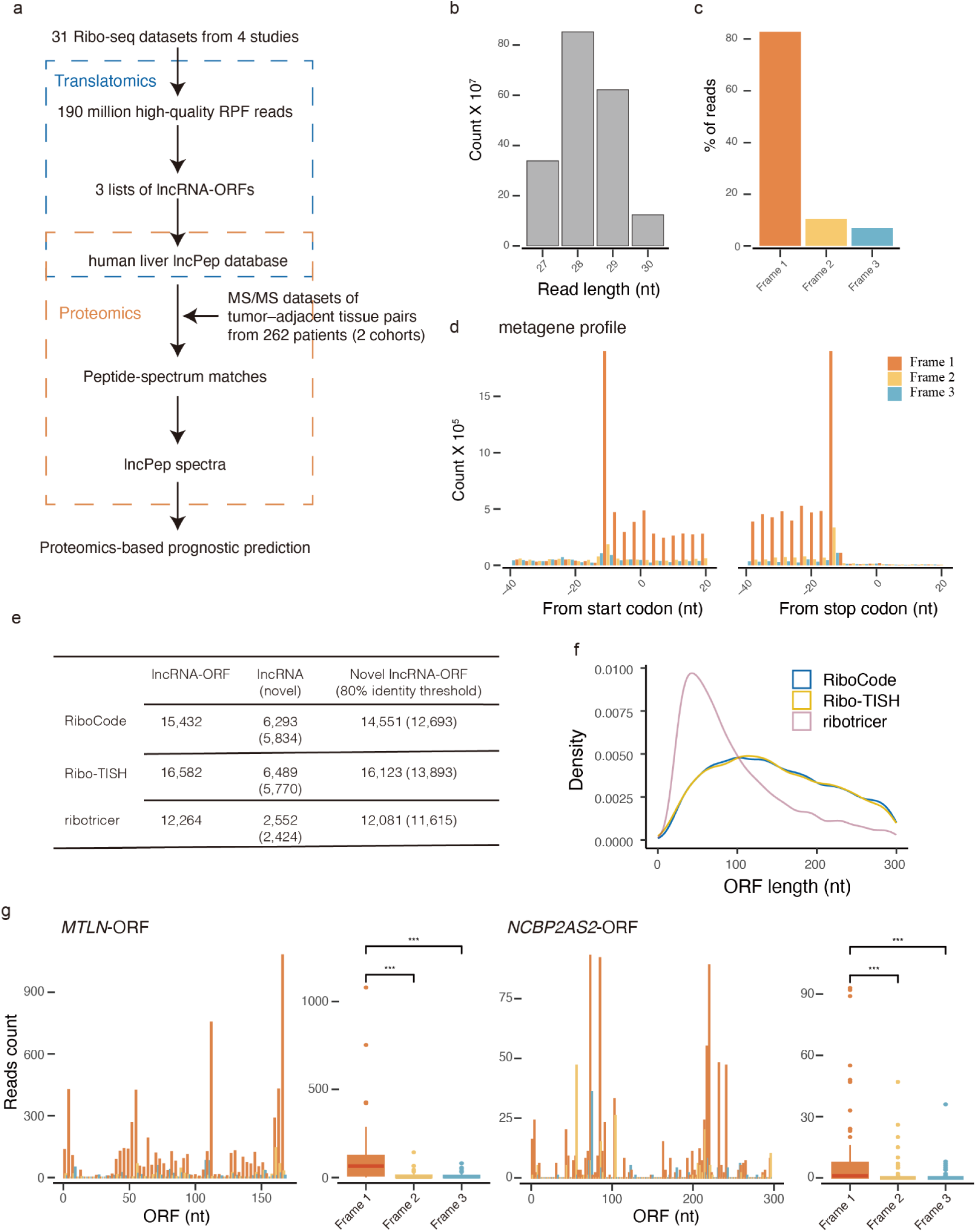
Identification of three lists of lncRNA-ORFs using high-quality Ribo-seq datasets and *de novo* ORF identifiers. a. Schematic flowchart of discovery analysis in this study. b. Bar chart demonstrating the overall reads distribution over 27-30 nt read lengths across Ribo-seq datasets. c. Overall frame reads distribution of the Ribo-seq datasets. d. Metagene profile showing triplet periodicity of P-site distribution in annotated ORFs. e. Summary of lncRNA-ORFs and parent lncRNAs predicted by RiboCode, Ribo-TISH, ribotricer *de novo* ORF identifiers. f. Kernel Density Estimation (KDE) showing length distribution of identified lncRNA-ORFs between 15 and 300 nt. g. P-site distribution showing triplet periodicity in *MTLN*-ORF and *NCBP2AS2*-ORF (Wilcoxon Rank Sum Test; ***, P < 0.001).

To systematically investigate coding potentials of putative lncRNA-ORFs in the human liver, we analyzed 31 ribosome profiling sequencing (Ribo-seq) datasets obtained from four independent studies deposited in the GEO database (73–76). These datasets were derived from human liver cell lines or tissues, including hepatocellular carcinoma (HCC) cell lines, untreated normal human hepatocytes, as well as cancerous and paracancerous tissues from HCC patients (Supplementary Table 1).

The processed Ribo-seq dataset comprised 190 million sequencing reads in total, with a length distribution ranging from 27 to 30 nucleotides (Figure 1b). Of the reads, 93.96% were mapped to annotated coding DNA sequence (CDS) regions of mRNAs, while the remaining reads were mapped to untranslated regions (UTRs) (4.53%) and ncRNA (1.52%) (Figure S1a). Triplet periodicity was confirmed within the annotated CDS, with over 80% of ribosome-protected fragments (RPF) reads aligned to frame 1 (Figure 1c). Additionally, these reads exhibited enrichment at both the CDS start and stop codons, with a consistent 12-nt offset (Figure 1d). Collectively, these metrics confirm the high quality of the Ribo-seq datasets used in this study (77).

To identify human liver active lncRNA-ORFs, we extracted candidate ORFs originating from lncRNAs annotated in GENCODE (version 46) and NONCODE (version 6) (37, 38), including GENCODE-annotated lncRNAs, GENCODE-annotated pseudogenes, and NONCODE-annotated lncRNAs. In addition to AUG, we included the near-cognate triplets GUG, CUG, and UUG as alternative start codons, as these codons have been shown to possess translational potential in previous studies (16). We assessed all candidate lncRNA-ORFs using three Ribo-seq-based active ORF identifiers: RiboCode, Ribo-TISH, and ribotricer (44, 45, 49).

For ORFs originating from different transcript isoforms of the same gene, or those sharing the same stop codon but with undefined start positions, we retained both the longest ORF and the optimal ORF when they were not identical. The optimal ORF was defined as the one with the lowest P-value from RiboCode or Ribo-TISH, or the highest phase score from ribotricer. Ribo-TISH identified 16,582 lncRNA-ORFs from 6,489 genes. RiboCode detected 15,432 lncRNA-ORFs from 6,293 genes, while ribotricer identified 12,264 lncRNA-ORFs from 2,552 genes (Figure 1e, Supplementary Table 2). The active ORFs identified by these methods were subsequently incorporated into downstream analyses.

The coding potentials of the identified lncRNA-ORFs were evaluated using CPC2, a machine learning-based tool that predicts the coding potential of the transcripts from which the ORFs originate (50). Among the 74,081 GENCODE-annotated transcripts, 8,400 were predicted to be translatable by at least one of the RiboCode, Ribo-TISH, or ribotricer identifier tools, with an average coding probability of 0.29. Additionally, 2,738 transcripts were predicted to be translatable by multiple tools, with an average coding probability of 0.39. For the 162,310 NONCODE-annotated transcripts, 17,860 were predicted to be translatable by at least one tool, with an average coding probability of 0.20, while 5,655 were predicted to be translatable by multiple tools, with an average coding probability of 0.29. Significantly higher coding probabilities based on sequence features of known mRNAs were observed for lncRNAs with ORFs detected by multiple identifiers and for those with ORFs detected by a single identifier, compared to lncRNAs without identified ORFs. Moreover, coding probabilities were higher for lncRNAs with ORFs detected by multiple identifiers than for those detected by a single identifier. These results indicate that the sequence features used for translation may have similarity between mRNAs and lncRNAs (Figure S1b, P < 0.01, Wilcoxon tests).

We compared the translated lncRNAs identified in this study with those reported in previous studies, including GENCODE Ribo-ORFs, UniProt lncRNA-encoded proteins/peptides, and the Chothani *et al*. 2022 study (51, 75, 78). At the gene level, RiboCode and Ribo-TISH showed higher concordance in their identified ORFs, whereas riboticer detected a larger number of active ORFs from fewer lncRNAs (Figure S1c). This difference may stem from riboticer’s use of the phase score rather than P-value-based algorithms, a feature previously reported to confer higher sensitivity to shorter ORFs (49). In line with this, the ORFs identified by ribotricer were significantly shorter than those predicted by RiboCode and Ribo-TISH (Figure 1f). Comparisons with published databases, both the GENCODE Ribo-ORF dataset and the Chothani *et al*. 2022 study showed large differences. Given that only human liver-derived cell or tissue data were used in this study, the lack of concordance may reflect differences in the translational landscapes of different tissues (Figure S1c). A similar trend to gene level comparison was observed at the lncRNA-ORF level (Figure S1d).

We compared the predicted lncRNA-ORFs with known CDS regions using three metrics: combined P-value (based on the distribution difference of RPF reads across frame 1, frame 2, and frame 3), phase score (ranging from 0 to 1, with higher scores indicating stronger triplet periodicity), and read density (RPF reads per ORF length). All three metrics showed comparable values between predicted lncRNA-ORFs and known CDSs, supporting the translational potential of these putative lncRNA-ORFs (Figure S1e).

Experimentally validated HCC-related lncRNA-ORFs, including KRASIM and MPM (23, 24), were successfully predicted by our Ribo-seq analysis by all three identifiers. We further examined the RPF tracks of two lncPeps: MPM (56 aa) and KRASIM (99 aa). Significant triplet periodicity pattern was observed in the 297 nt and 168 nt ORFs of lncRNA *MTLN* (P=1.09e-15, Stouffer’s Method), and lncRNA *NCBP2AS2* (P=6.17e-14, Stouffer’s Method) (Figure 1g, Figure S1f). These results provide additional validations for the translational potential of the lncRNA-ORFs identified in this study.

### Identification of lncPeps from mass spectrometry data

We constructed three proteogenomics candidate databases from the identified lncRNA-ORFs. Database 1 included lncPeps predicted by two or more of the three tools, totaling 9,451 lncPeps (RiboCode ∩ Ribo-TISH) ∪ (RiboCode ∩ ribotricer) ∪ (Ribo-TISH∩ ribotricer). Database 2 included lncPeps predicted by at least one tool, totaling 33,083 lncPeps (RiboCode ∪ Ribo-TISH ∪ ribotricer). Database 3 included all candidate ORFs from transcripts showing RPF signals. In this case, no triplet periodicity filtering was applied, resulting in 288,383 ORFs (Figure 2a).

**Fig 2:**
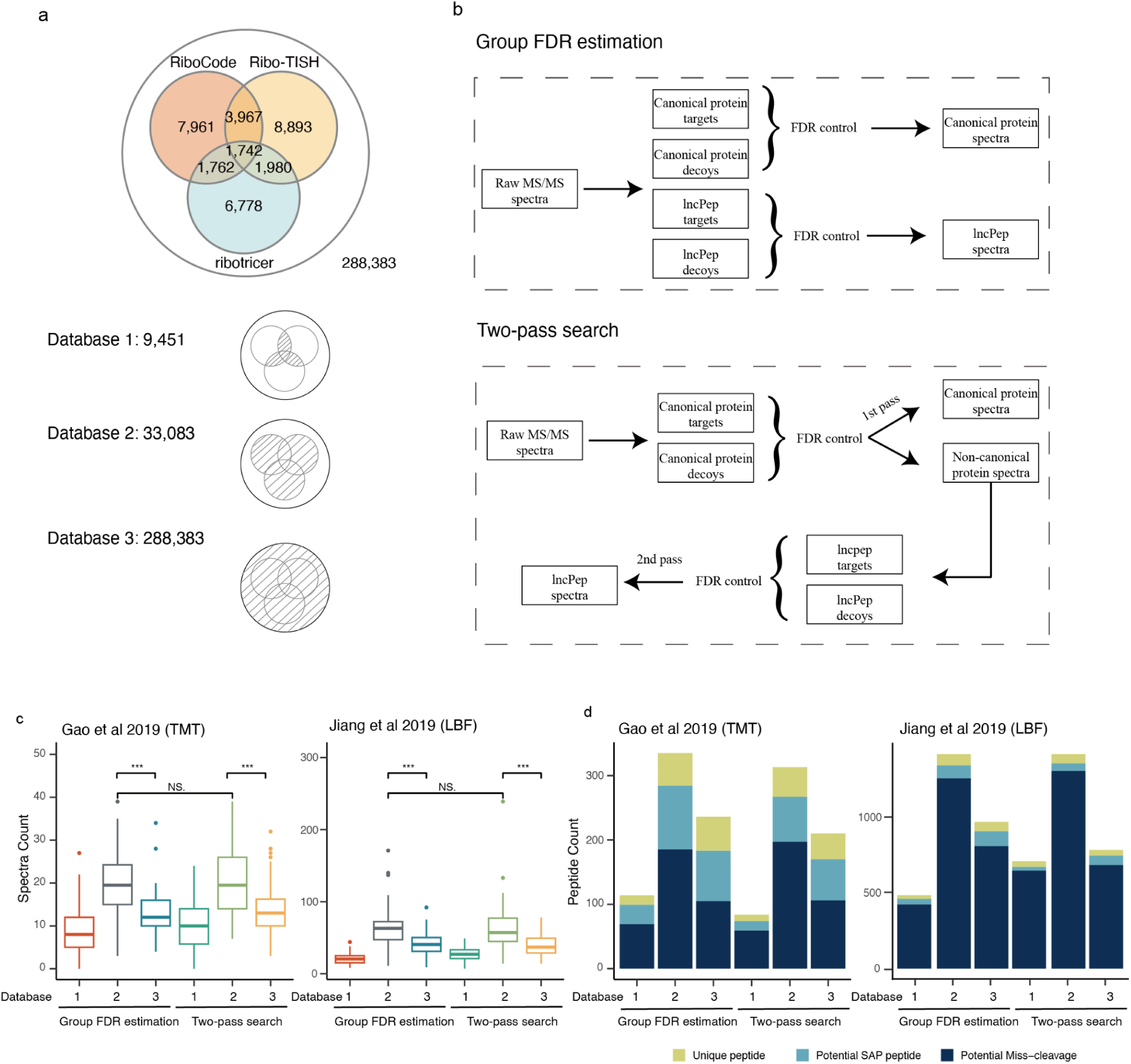
PSM group FDR control for optimizing the integrated lncPep database. a. Venn diagram showing the three overlapping lncRNA-ORF lists and potential proteogenomic lncPep databases translated from the lncRNA-ORFs (database 1: lncRNA-ORFs detected in at least two identifiers; database 2: lncRNA-ORFs detected in at least one identifier; database 3: all potential lncRNA-ORFs). b. Flowchart of the Group FDR estimation and the Two-pass search as statistical strategies in selecting lncPep databases (FDR threshold 1%). c. Box-and-line plots demonstrating the number of putative lncPep-sourced MS/MS spectra identified using each lncPep database: Gao *et al*. (2019) (left) and Jiang *et al*. (2019) (right). d. Bar charts showing the number of unique peptides, potential SAP peptides, and potential miss-cleaved peptides detected; Gao *et al*. (2019) (left) and Jiang *et al*. (2019) (right).

lncRNA-encoded peptides differ from canonical proteins in length, abundance, and sequence features, which can affect the accuracy of peptide identification in computational proteomics analyses. Therefore, group-specific false discovery rate (FDR) estimation for lncPep-sourced spectra is critical to ensure reliable peptide identification (79, 80). We compared the performance of two FDR control strategies: group-specific FDR estimation and the two-pass search method for identifying lncRNA-derived peptide spectra (56, 57). The group-specific FDR estimation approach identifies peptides from both canonical proteins and lncRNA-encoded peptides but applies FDR control separately for each group. In contrast, the two-pass search strategy first performs peptide identification and FDR control on canonical proteins and then evaluates whether the spectra not attributed to canonical proteins may originate from lncRNA-encoded peptides (Figure 2b).

To evaluate the performance of the two FDR control methods and the three databases, we analyzed two test datasets using the FragPipe platform. Each test dataset consisted of cancerous and paracancerous tissue samples from five patients, randomly selected from the larger cohorts reported by Jiang *et al*. and Gao *et al* (6, 36).

Database 2, which has the intermediate stringency between database 1 (most stringent) and database 3 (most permissive), outperformed database 1 and database 3 in identifying higher numbers of lncPep-sourced spectra (Figure 2c). Notably, the higher detection sensitivity of database 2 suggests that Ribo-seq-guided proteogenomic approaches impact downstream proteomics analyses. A comparison of Group FDR estimation and two-pass search method found no significant difference in the number of identified lncPep-sourced spectra between the two strategies (Figure 2c).

Peptide fragments fully aligned with canonical proteins but lacking a trypsin cleavage site were considered potential trypsin miscleavages—a common occurrence in trypsin-based proteomics—and were filtered out from subsequent analyses (81). Single amino acid polymorphisms (SAPs) frequently occur in peptides. Therefore, we applied an additional filtering step to spectra that are potentially originating from lncPeps. Fragments aligned to canonical proteins with a single amino acid mismatch were regarded as a potential SAP product. Peptide fragments that could not be aligned to any canonical proteins, after allowing one mismatch, were considered unique and retained for subsequent lncPep identification and quantification. A substantial number of lncPep spectra were mapped to missed cleavage or SAP events in both datasets during the above filtering process (Figure 2d).

Based on these results, we finalized the rest of the analysis steps for lncPep discovery, which involved performing proteomics analysis using database 2 and controlling the false discovery rate (1% FDR threshold) with group-specific estimation. The detailed workflow is illustrated in Figure S2.

### Physicochemical properties limited the detection of lncPep by mass spectrometry

We analyzed the full proteomic datasets from two previously reported cohorts. In the original studies, Jiang *et al*. profiled 103 hepatocellular carcinoma (HCC) patients at clinically early stages, whereas Gao *et al*. reported 159 HCC patients associated with hepatitis B virus infection (6, 36). By integrating the two datasets, we identified a total of 105 lncPep groups originating from 165 potential lncRNA-ORFs (Supplementary Table 3). lncPeps that are ambiguous with respect to their ORF start codon or source transcript isoform, or cannot be distinguished based on proteomic data, were grouped into a single lncPep group.

From the 105 lncPeps discovered, we present three representative lncPeps nested in: *PPIAP79* lncORF1, *POTEKP* lncORF2, and *HNRNPA1P36* lncORF1, along with their corresponding RPF tracks and representative MS spectra (Figure 3). *PPIAP79* lncORF1 encodes a 92aa small peptide, from which two unique peptide fragments were detected. *POTEKP* lncORF2 encodes a 419aa peptide that has been cataloged in UniProt (82). *HNRNPA1P36* lncORF1 encodes a peptide comprising 275 amino acids.

**Fig 3:**
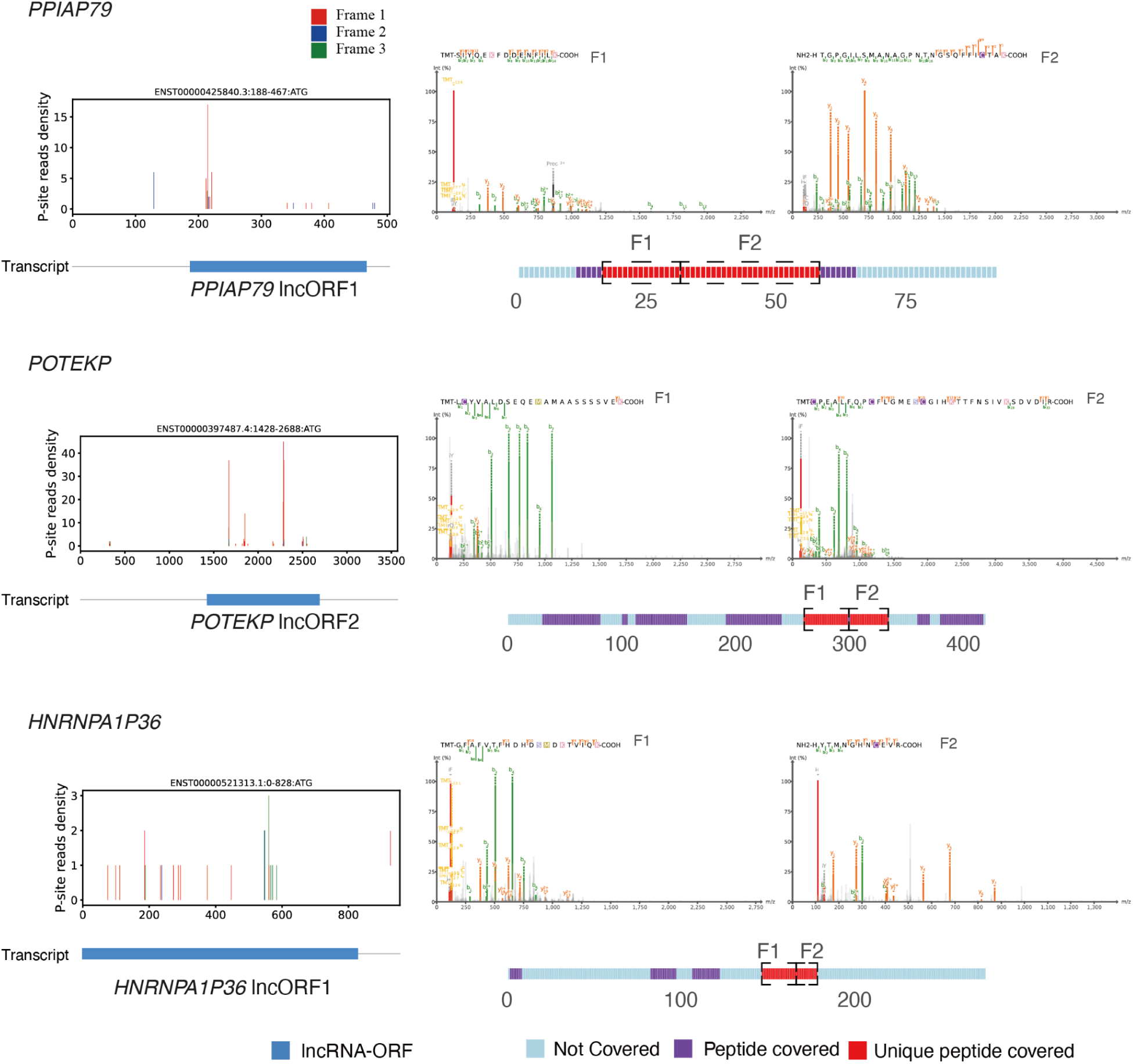
RPF tracks and matched mass spectra of three lncPep examples. RPF tracks (left) and representative unique peptide MS spectra (right) of *PPIAP79* lncORF1 (top), *POTEKP* lncORF2 (middle), and *HNRNPA1P36* lncORF1 (bottom).

We traced the sources of these lncPeps and found that 84.8% were exclusively identified from the newly generated lncPeps discovered in this study. When comparing specific ORF detection algorithms, most lncPep detections were attributed to the overlapping lncRNA-ORF pools from RiboCode and Ribo-TISH. Some lncPeps were detected exclusively from the RiboCode lncRNA-ORF pool, while only five ribotricer-predicted ORFs were detected in the mass spectrometry data. In terms of gene origins, most lncPeps were derived from GENCODE-annotated pseudogenes, with some originating from NONCODE-annotated lncRNA genes, and a few from GENCODE-annotated lncRNA genes (Figure 4a).

**Fig 4:**
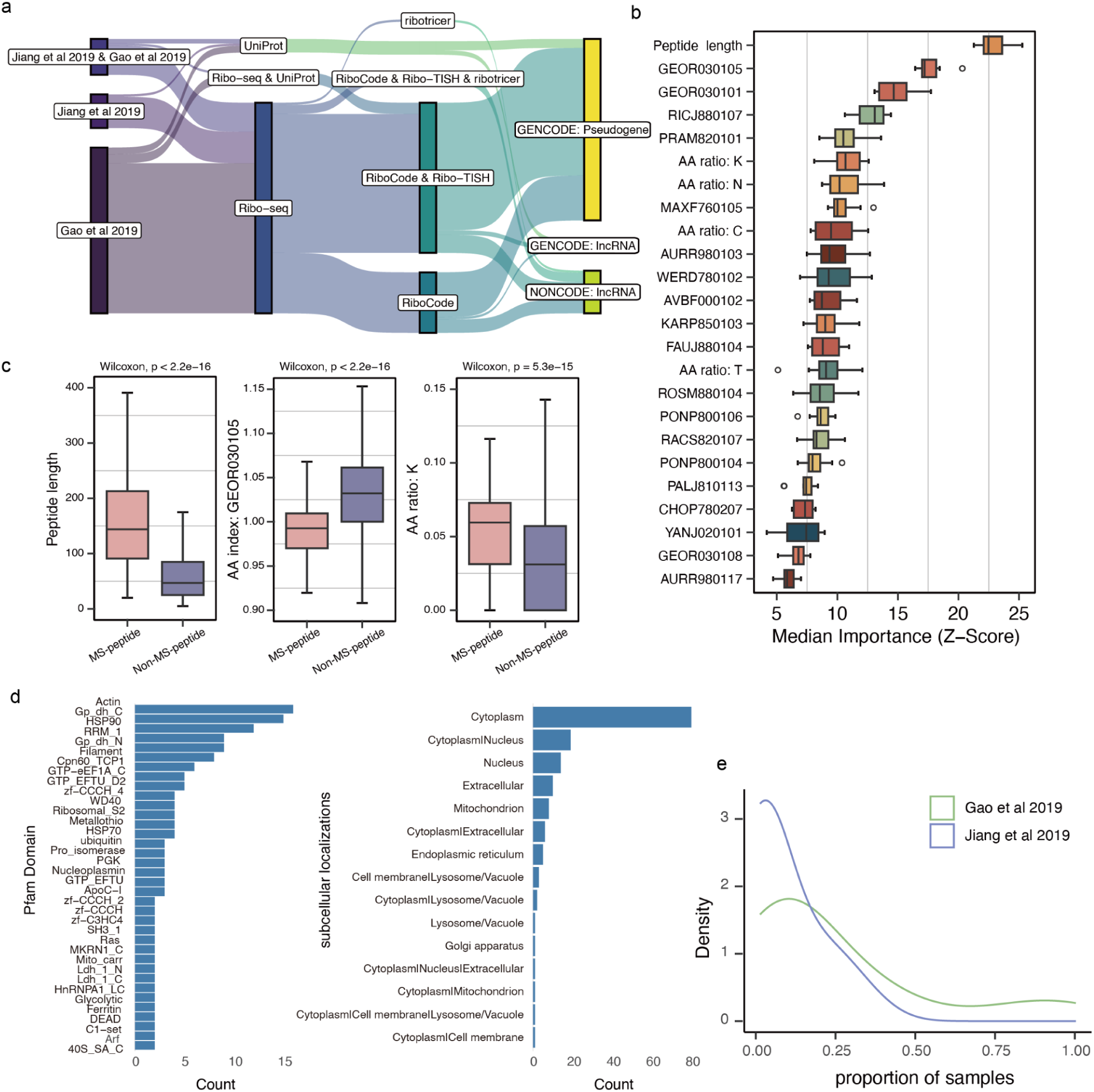
lncPeps detected in HCC patient proteomic datasets. a. Sankey diagram visualizing traffic flow of the identified lncPeps between origin of datasets, databases, ORF identifiers, and biotypes annotated in GENECODE and NONCODE projects. b. Box plots displaying the Importance of peptide physicochemical properties ranked by Boruta’s algorithm (higher Importance indicates stronger contribution to distinguishing MS-peptides from non-MS-peptides). c. Box plots showing peptide length (left), GEOR030105 score (center), and lysine ratio (right) in MS-peptide group and non-MS-peptide group. Statistical p values are from Wilcoxon Rank Sum Test. d. Count of Pfam domains and predicted subcellular localizations of the detected lncPeps. e. Detection rates of lncPeps across samples.

To understand why a large number of lncRNA-ORFs were not matched by mass spectra signals, we investigated the physicochemical properties of lncPeps derived from Ribo-seq data that have been detected by proteomic mass spectrometry (MS-peptides) and lncPeps derived from Ribo-seq data but undetected by proteomic mass spectrometry (Non-MS peptides). The considered properties included 577 features, such as peptide length, molecular weight, theoretical isoelectric point, individual amino acid ratios, amino acid family ratios, and AAindex features. After filtering these features, we used LASSO regression for modeling and retained 24 key features that make significant contributions to the distinction.

By analyzing the average Importance of variables in each feature, we identified peptide length as the most influential feature. Notably, MS-peptides exhibited significantly longer sequences compared to Non-MS peptides (Figure 4b-c). This observation also explains why we detected less ribotricer-predicted lncORFs matched by MS spectra, as the ribotricer algorithm favors shorter ORFs (Figure 1f). Additionally, two AAindex features, GEOR030105 and GEOR030101, showed strong correlations, both associated with the presence of protein domain linkers in peptide structures (Figure 4b-c) (83). MS-peptides exhibited a significantly lower propensity for domain linkers. Furthermore, we observed that the AAindex feature RICJ880107, which reflects a tendency for alpha-helix formation, had a significantly lower presence in MS-peptides (84).

We performed machine learning modeling using RandomForest algorithm based on these physicochemical features. For the complete set of lncRNA-ORF, we used 70% of the peptides as a training set and the remaining 30% as a test set. Peptide length, amino acid composition, and AAindex features were each modeled separately using the 24 features selected by LASSO. The AUC values for these models in the test set were 0.8433, 0.7995, and 0.9364, respectively. When modeling was performed using all selected features combined, the AUC value increased to 0.9547 (Figure S3). These results indicate that there are physicochemical differences between MS-peptides and Non-MS-peptides, which may explain why some lncPeps are more difficult to detect using the trypsin-based, MS proteomic technology employed in the two patient studies.

To gain insights into the function of the discovered lncPeps and the cellular pathways they impact, we performed Pfam domain analysis. The most frequently occurring domains were the Actin family, Gp_dh_C, and HSP90 family (Figure 4d). Subcellular localization predicted by DeepLoc2 predominantly localized the lncPeps to the Cytoplasm and Nucleus regions, with smaller fractions assigned to other compartments (e.g., mitochondria, ER, extracellular) (Figure 4d).

Finally, we calculated the detection rates of these lncPeps across patient samples. In the Jiang *et al*. 2019 dataset, the average detection frequency of lncPeps was 0.085, indicating that on average a lncPep was detected in 8.5% of the samples. In contrast, the Gao *et al*. 2019 dataset, which utilized TMT technology, showed a higher detection rate for lncPeps, with an average frequency of 0.25 (Figure 4e).

### Novel lncPeps show potential as prognostic biomarkers for HCC patients

We performed abundance estimation and biostatistical analysis of these lncPeps to assess their potential clinical utility. PSMs corresponding to unique peptide fragments were used for quantification, with MSstats and MSstatsTMT applied to the LBF and TMT datasets, respectively (60, 61).

We first performed principal component analysis (PCA) on the proteomics expression matrix. In the Jiang *et al*. dataset, PCA of the Merged expression (canonical proteins and lncPep expression) revealed a clear separation between tumor and non-tumor tissues, whereas no such separation was observed with lncPep expression alone. Similarly, partial least squares discriminant analysis (PLS-DA), a supervised learning approach, failed to discriminate tumor from non-tumor tissues based solely on lncPep expression (Figure S4a-c). This may be attributed to the sparseness of lncPeps expression matrix, which limits their biostatistical significance (Figure 4e). Batch correction was confirmed in the Gao *et al*. dataset. Subsequent analyses demonstrated that tumor and non-tumor tissues were best separable at the Merged expression level. PCA of lncPep expression captured partial group separation, which was further enhanced by PLS-DA, suggesting that lncPep expression harbors discriminatory information (Figure S4d-g).

To further explore the potential of lncPeps as biomarkers for HCC, we constructed predictive models based on the three types of input data (canonical protein data, lncPep data, or merge of the two) and three distinct tasks comprising tumor classification, survival prognosis, and recurrence prognosis.

In the tumor tissue classification models using the Jiang *et al*. 2019 dataset, the lncPep-based model performed poorly, and incorporating lncPep expression did not significantly improve the performance of the canonical protein-based model (Figure 5a, lncPep model AUC = 0.56; likelihood ratio test for the merge model vs. the canonical protein model, P = 0.68). Similar trends were observed in the Gao *et al*. 2019 dataset, where the lncPep-based model achieved an AUC of 0.95 but remained inferior to the protein-based model, and adding lncPep expression did not significantly improve performance compared with using canonical protein expression alone (Figure 5a, likelihood ratio test for the merge model vs. the canonical protein model, P = 1).

**Fig 5:**
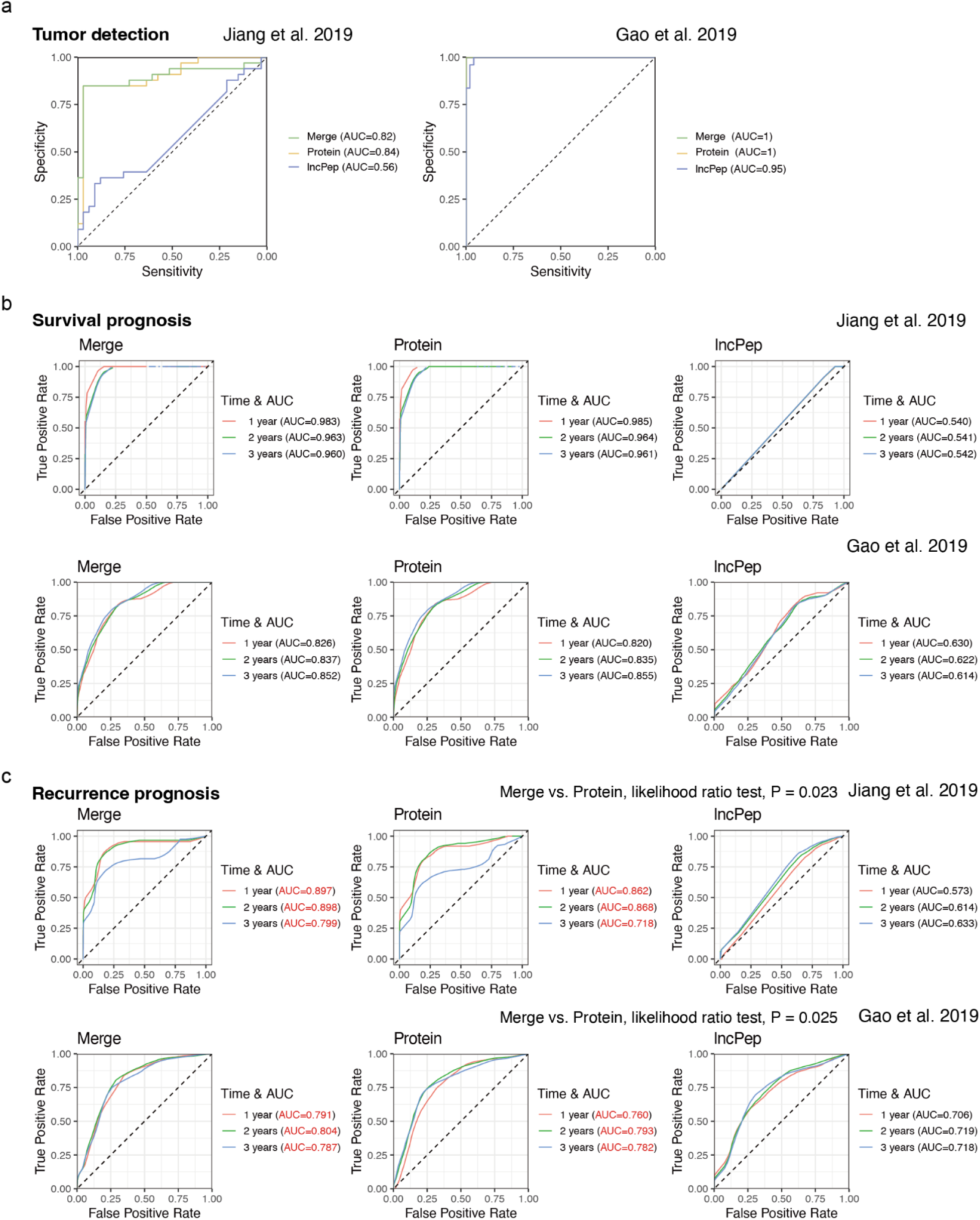
Improved performance of the HCC recurrence prediction model by lncPep expression information. AUC curves of models based on canonical protein expression, lncPep expression, or merged for a. tumor detection, b. survival prognosis, and c. recurrence prognosis. Red color denotes AUC values between significantly different Merge and Protein models detected by Likelihood Ratio Test.

For survival prognosis models, incorporating lncPep expression did not improve the performance of the canonical protein-based model in either the Jiang *et al*. 2019 or Gao *et al*. 2019 datasets (Figure 5b; likelihood ratio test for the merge model vs. the canonical protein model, P = 0.92 for Jiang *et al*. 2019 and P = 0.082 for Gao *et al*. 2019 datasets). In contrast, for recurrence prognosis models, inclusion of lncPep expression significantly enhanced predictive performance in both datasets (Figure 5c; likelihood ratio test for the merge model vs. the canonical protein model, P = 0.023 for Jiang *et al*. 2019 and P = 0.025 for Gao *et al*. 2019 datasets). These results indicate functional relevance and thus positive contribution of lncPeps to prognostic biomarker development for HCC.

### Individual lncPeps demonstrate correlations with tumors and patient prognosis

We then investigated the differences in individual lncPep expression between tumor and non-tumor tissues, as well as the correlation between lncPep expression and patient survival and recurrence rates, using logistic regression and Cox regression analyses. To address missing values in the lncPep expression matrix, we employed two approaches: filtering out samples with missing data or applying minimal value imputation. In addition, we analyzed the TCGA-LIHC (The Cancer Genome Atlas Liver Hepatocellular Carcinoma Collection) dataset to explore the impact of expression changes in the RNAs encoding these lncPep (85).

Through this analysis, we identified a series of lncPeps whose expression was associated with cancerous tissue or associated with patient prognosis (Figure 6a). We first focused on peptides that exhibited consistent effects across both the Jiang *et al*. and Gao *et al*. datasets. lncPep *PPIAP79* lncORF1 demonstrated a significant association with tumor tissues in the Jiang *et al.* dataset (logistic regression, P = 0.0013 for remove missing values, P = 0.13 for minimum value imputation) and also showed a significant correlation with tumor tissues in the Gao *et al*. dataset using both methods (Figure 6b, logistic regression, P = 0.033 for remove missing values, P = 0.021 for minimum value imputation). Differential expression analysis revealed that *PPIAP79* lncORF1 was significantly upregulated in cancerous tissues in the Jiang *et al*. dataset (Wilcoxon test, P value = 8e-7 for remove missing values, P = 0.13 for minimum value imputation) and in both methods using the Gao *et al*. dataset (Figure 6b, Figure S5a, Wilcoxon test, P value = 0.017 for remove missing values, P = 0.01 for minimum value imputation). Similarly, *SEPTIN7P8* lncORF1 showed a significant correlation with non-tumor tissues in the Jiang *et al*. dataset (logistic regression, P = 0.94 for remove missing values, P = 0.045 for minimum value imputation) and in both methods of the Gao *et al*. dataset (Figure 6b, logistic regression, P = 0.023 for remove missing values, P = 0.023 for minimum value imputation). Differential expression analysis indicated significant downregulation in paracancerous tissues in the Jiang *et al*. dataset (Wilcoxon test, P value = 0.7 for remove missing values, P = 0.022 for minimum value imputation) and in the Gao *et al*. dataset (Wilcoxon test, P value = 0.033 for remove missing values, P = 0.012 for minimum value imputation) (Figure 6b, Figure S5b). We were unable to compare RNA expression levels of these pseudogenes in the TCGA-LIHC dataset due to insufficient data (Figure 6a).

**Fig 6:**
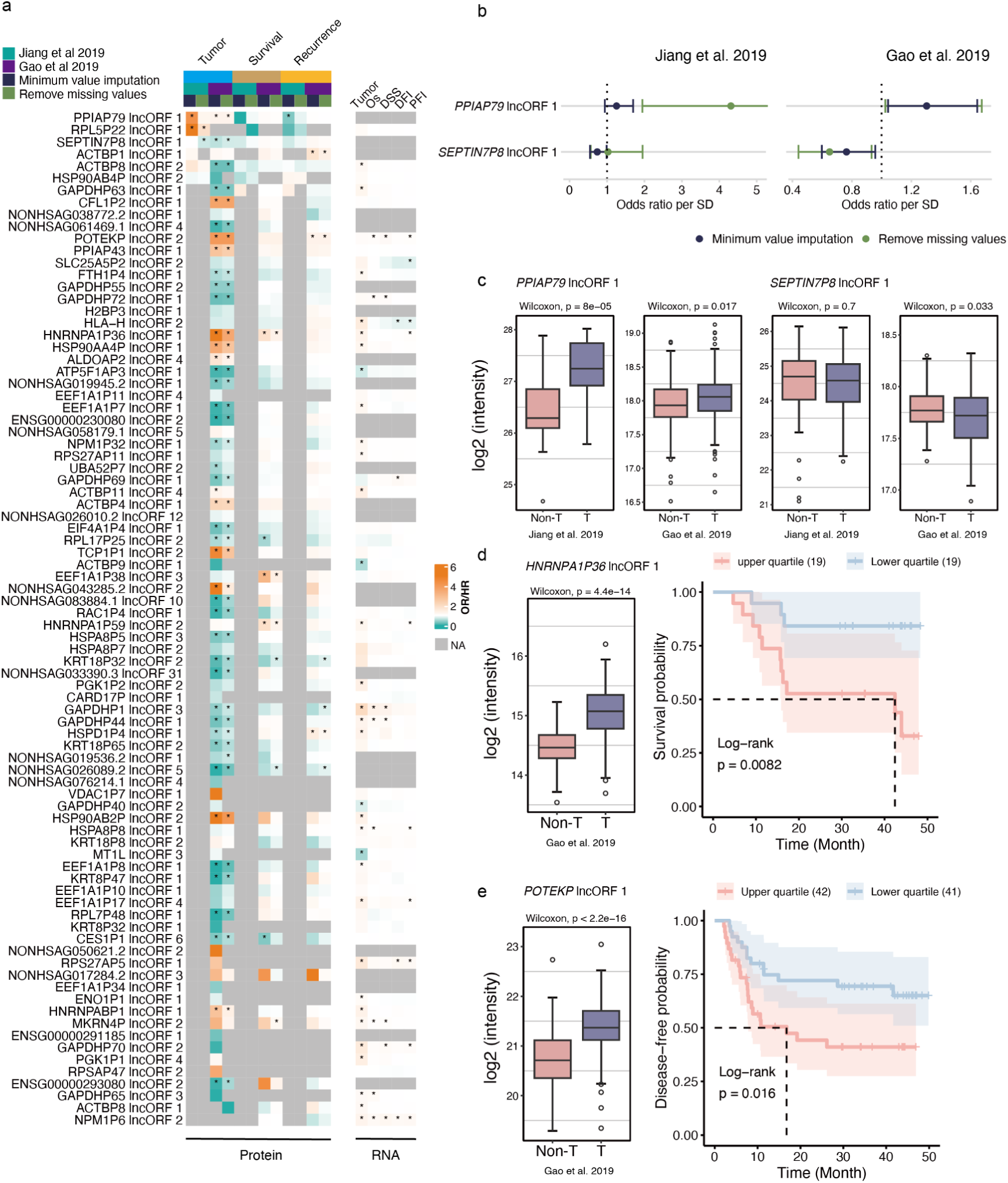
Newly discovered lncPeps are associated with tumor, patient survival, and recurrence. a. Heatmap shows the odds ratio (OR) for tumor, hazard ratio (HR) for overall survival prognosis, and HR for recurrence prognosis, mapped against each newly discovered lncPep (left) and sourced lncRNA (right). Statistical significance was assessed using logistic regression or Cox regression; *, P < 0.05. OS, Overall Survival; DSS, Disease-Specific Survival; DFI, Disease-Free Interval; PFI, Progression-Free Interval. b. Forest plots showing the association between expression of lncPeps (*PPIAP79* lncORF1 and *SEPTIN7P8* lncORF1) and OR for tumor in Jiang *et al*. 2019 and Gao *et al*. 2019. c. Box plots showing expression of *PPIAP79* lncORF1 and *SEPTIN7P8* lncORF1 in non-tumor tissue (Non-T) and tumor tissue (T). d. Box plot showing expression of *HNRNPA1P36* lncORF1 in non-tumor and tumor tissues, and Kaplan–Meier (KM) analysis of overall survival probability in patients stratified by HNRNPA1P36 lncORF1 expression (upper vs. lower quartile). e. Box plot showing expression of *POTEKP* lncORF2 in Non-tumor and tumor tissues, and KM analysis of recurrence prognosis. Statistical significance was assessed using Log-rank test.

We also detected lncPeps with significant associations with patient prognosis. For example, *HNRNPA1P36* lncORF1 was upregulated in tumor tissues and significantly correlated with patient survival (Figure 6d, Figure S5c, assessed by both imputation methods). Furthermore, Cox regression and log-rank tests confirmed that its expression is associated with survival outcomes. Similarly, *POTEKP* lncORF2 was significantly upregulated in tumor tissues (Figure 6e, Figure S5d, assessed by both imputation methods); Cox regression and log-rank tests confirmed its association with patient recurrence outcomes (Figure 6e, Figure S5d). Analysis of TCGA transcriptomic data showed that, although *POTEKP* RNA levels were not differential between tumor and non-tumor tissues, they were significantly associated with patient prognosis, specifically overall survival (OS), disease-specific survival (DSS), and progression-free interval (PFI) (Figure S6a; assessed by both Cox regression and log-rank tests). In contrast, *HNRNPA1P36* RNA was significantly upregulated in tumor tissues, consistent with the protein-level data, and its expression correlated with PFI-associated prognosis (Cox test P = 0.03, log-rank test P = 0.053) (Figure S6b).

In order to explore the biological significance of the three lncPeps, canonical proteins were ranked according to their Pearson correlation with lncPep expression and then subjected to GSEA. The results indicated that *PPIAP79* lncORF1 expression was positively correlated with proteins involved in mitochondrial gene expression and mitochondrial translation. *POTEKP* lncORF2 expression showed significant positive correlations with proteins related to the humoral immune response, complement activation, and other immune-related processes. In contrast, *HNRNPA1P36* lncORF1 expression was positively correlated with proteins involved in mRNA processing, chromatin organization, and related cellular processes (Figure S7).

Collectively, these results demonstrate that specific lncPeps are associated with HCC tumors or patient prognosis, and for particular lncPeps, the associations can also be observed at the transcript level. These findings suggest that lncPeps may have functional roles in HCC tumorigenesis and progression, or alternatively, represent byproducts of these processes.

## Discussion

Proteogenomics, defined as the integration of proteomics with genomics and/or transcriptomics, has facilitated the annotation of an increasing number of lncPeps. However, this approach still faces several limitations. Specifically, the sequence content of proteogenomic databases serves as an informed prior about sample composition, and incorrect prior assumptions can negatively impact peptide and protein identification. In practice, this means that larger reference database search space reduces identification sensitivity at a given false discovery rate (86, 87). To overcome these limitations, we leveraged recent advances in translatomics from tissue-specific ribosome profiling experiments. Our strategy begins with the prediction of lncRNA-derived ORFs using Ribo-seq data to construct a tailored proteogenomic database. MS-based proteomics data are then mapped to this database, and novel lncPeps are identified following rigorous filtering steps. This approach enables both stringent peptide detection and quantitative comparisons across patients. Using this strategy, we identified a total of 105 lncPep groups from the cancerous and paracancerous tissues of two independent HCC cohorts.

To optimize the computational proteomics pipeline for maximizing the detection of lncPep-derived spectra under the same FDR control level (1%), we evaluated two group FDR control methods and three different tolerance levels of the lncRNA-ORF database. Our results showed that, although the two-pass search method identified substantially more spectra matching lncPeps compared to group FDR estimation. However, the sensitivity for lncPep-derived spectra was comparable between the two methods. For different levels of database tolerance, we found that the moderately permissive database 2 exhibited higher sensitivity. This observation is consistent with the previously reported notion that a reference database that is too small may fail to match spectra, whereas an excessively large database reduces identification sensitivity. Incorporating Ribo-seq–derived prior information played a key role in improving the accurate identification of these lncPeps.

Of note, among the lncPeps identified in this study, the majority originated from pseudogene-derived lncRNAs rather than from intergenic or antisense lncRNAs. This observation is consistent with previous reports in human epidermal carcinoma and B-lymphoblast cell lines (88, 89). Pseudogenes are duplicated copies of protein-coding genes that have lost the ability to produce functional protein products identical to their parental genes, while retaining a high degree of sequence similarity. Based on this property, pseudogenes have long been hypothesized to exert regulatory functions through the transcription of lncRNAs that modulate the expression of their parental genes (90). Notably, pseudogenes often exhibit cancer-specific expression patterns (91), and in some cases their transcripts are more abundant in tumor tissues than in normal counterparts (92, 93). In contrast, previous immunopeptidome datasets have identified a substantially greater number of lncPeps derived from intergenic or antisense lncRNAs than from pseudogenes (89). This distinction may reflect that lncPeps encoded by lncRNAs of different genomic origins fulfill distinct biological roles.

The results of this study imply that physicochemical properties are key determinants of whether or not lncRNA-encoded peptides can be detected by protein mass spectrometry, and in particular that ORF length is the most important feature, which is in agreement with Yang et.al in their studies of lncRNA- and mRNA-derived microproteins, where logistic regression model showed that ORF length is the most important feature in determining whether microproteins can be detected in mass spec(94). The length may influence whether these non-canonical peptides are stabilized in cells and/or whether they can be hydrolyzed by trypsin. This length bias may render the majority of ORFs predicted by ribotricer undetectable by proteomic mass spectrometry. Both in this study and in previous reports, ribotricer showed higher sensitivity toward short ORFs. In addition, we found that these MS-peptides are less likely to constitute protein linker regions, which are typically enriched in intrinsically disorder residues (95).

Through biostatistical analyses, we identified a series of lncPeps associated with tumor tissues or patient prognosis, among which we focused on three peptides: *PPIAP79* lncORF1, *POTEKP* lncORF2, and *HNRNPA1P36* lncORF1. Of these, *POTEKP* lncORF2 has been cataloged in UniProt, whereas the other two were discovered in this study. Both *POTEKP* lncORF2 and *HNRNPA1P36* lncORF1 were differentially expressed between tumor and non-tumor pairs and show significant associations with patient prognosis. Interestingly the correlation with prognosis was also observed for RNA expression in the TGCA RNA-seq data, thus the noncanonical protein coding may provide an alternative mechanism in addition to competing endogenous RNA (ceRNA) to explain the correlation between cancer and RNA expression. Previous reports have suggested a *POTEKP*-encoded product showing sequence similarity to beta-actin and has implications in hepatocellular carcinoma, consistent with our findings (82). In this study, the observed association between *POTEKP* lncORF2 and patient prognosis may reflect a potential role of actin cytoskeleton components in HCC progression. Nevertheless, our understanding of the *POTEKP* remains limited.

We noticed that none of the three lncRNA-ORFs demonstrated strong RPF tracks in the Ribo-seq analysis. This phenomenon may be due to the fact that they are all derived from pseudogenes that have similar sequences to the corresponding protein-coding genes, causing mapping ambiguity of the RPF reads, especially considering that the lengths of the RPFs are short.

The results collectively affirm the discovery potential of the proteogenomic pipeline constructed in this study. Although our understanding of these non-canonical peptides remains highly limited, accumulating evidence suggests that they play important biological roles in human physiological functions and diseases. Therefore, our work warrants future investigations of lncPeps in more cancer types to help better characterize tumor heterogeneity and expand the pool of biomarkers, and ultimately benefit patients.

## Data availability

This study utilized publicly accessible datasets, with the Ribo-seq data listed in Supplementary Data 1, and protein mass spectrometry data obtained from the iProX and CPTAC projects. The R scripts for summarizing the results and generating the figures are available at https://github.com/lbwfff/lncRNA_derived-peptide_in_HCC.

## Supplemental data

This article contains supplemental data.

## Author contributions

Bingwu Li: Conceptualization; Formal analysis; Methodology; Data presentation; Writing original draft, reviewing & editing. Kandarp Joshi: Conceptualization; Reviewing and editing draft. Dan Ohtan Wang: Conceptualization; Writing original draft, review & editing; Funding acquisition; Supervision.

## Acknowledgments

The authors would like to thank all lab members in the RNA-MIND Lab at NYUAD for helpful discussions. This work is supported by the Research Award from NYUAD to DOW. This research was carried out on the High Performance Computing resources at New York University Abu Dhabi.

## Abbreviations

HCC: hepatocellular carcinoma
lncRNA: long noncoding RNA
lncPep: lncRNA-encoded peptide
ORF: open reading frames
sORF: small ORF
lncRNA-ORF: long non-coding RNA ORF
Ribo-seq: ribosome profiling sequencing
CDS: coding sequence
UTR: untranslated region
ncRNA: non-coding RNA
PSM: peptide spectrum match
nsSNP: non-synonymous single nucleotide polymorphism
ROS: reactive oxygen species
GSEA: gene set enrichment analysis
FDR: false discovery rate.

---

**Figure S1:**
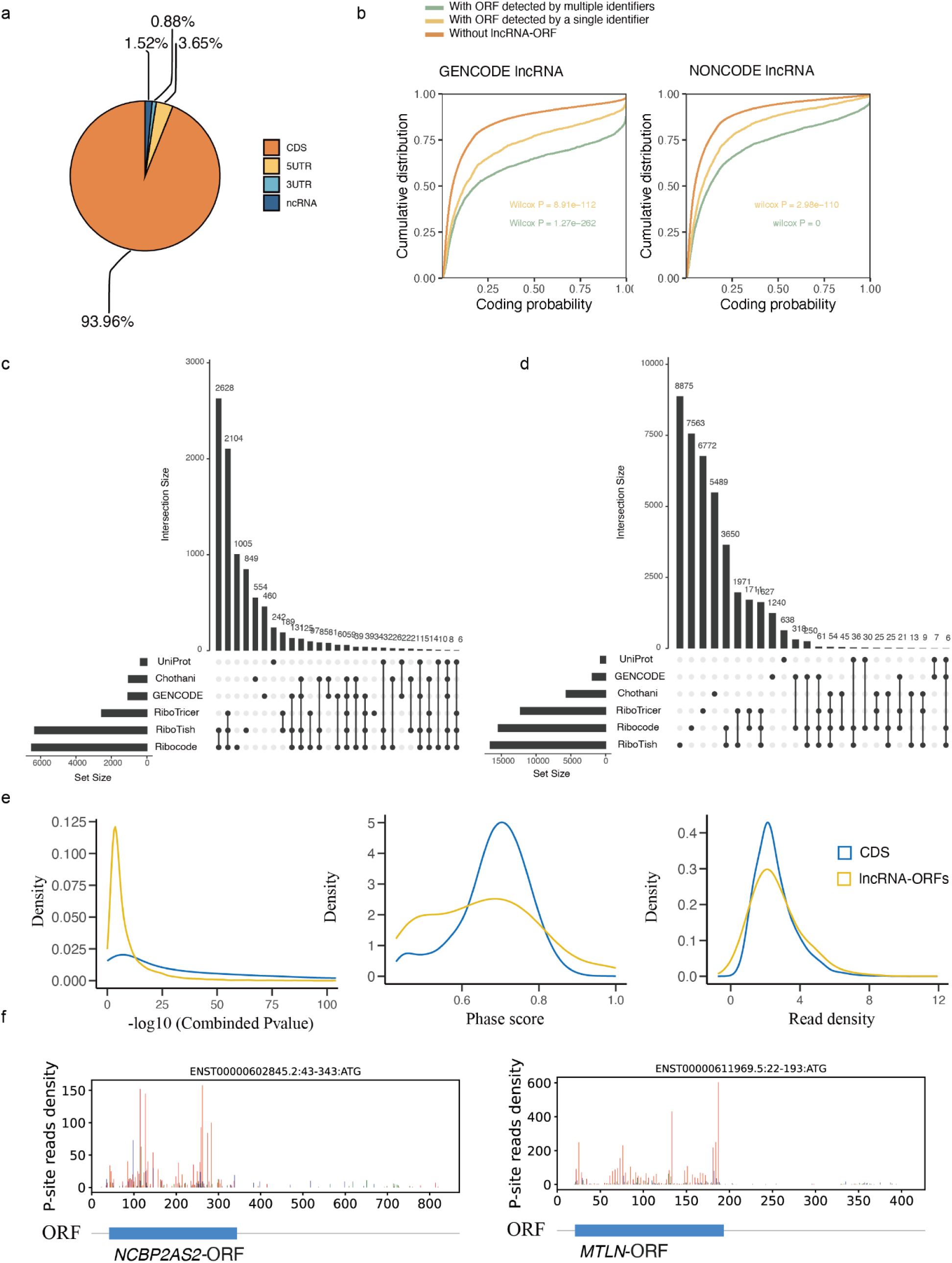
Supplementary figure for Figure 1. a. Distribution of Ribo-seq reads across transcriptomic features, including coding sequences (CDS), untranslated regions (5UTR, 3UTR), and non-coding RNAs (ncRNAs). b. Cumulative distribution of CPC2-predicted coding scores for three categories of lncRNA transcripts: transcripts lacking predicted ORFs, transcripts with ORFs predicted by a single tool, and transcripts with ORFs predicted by multiple tools. Left: lncRNAs annotated in GENCODE; right: lncRNAs annotated in NONCODE. c–d. Overlap of identified lncRNA genes (c) and lncRNA-ORFs (d) detected by three tools in this study, with annotations from GENCODE, Chothani et al. (2022), and the UniProt database. Only bars representing counts greater than 5 are shown. e. Distribution of combined P-values (left), phase scores (middle, calculated by ribotricer), and read densities (right) for predicted lncRNA-ORFs and known coding sequences (CDSs). f. Distribution of RPF reads across transcripts of *NCBP2AS2* and *MTLN*.

**Figure S2:**
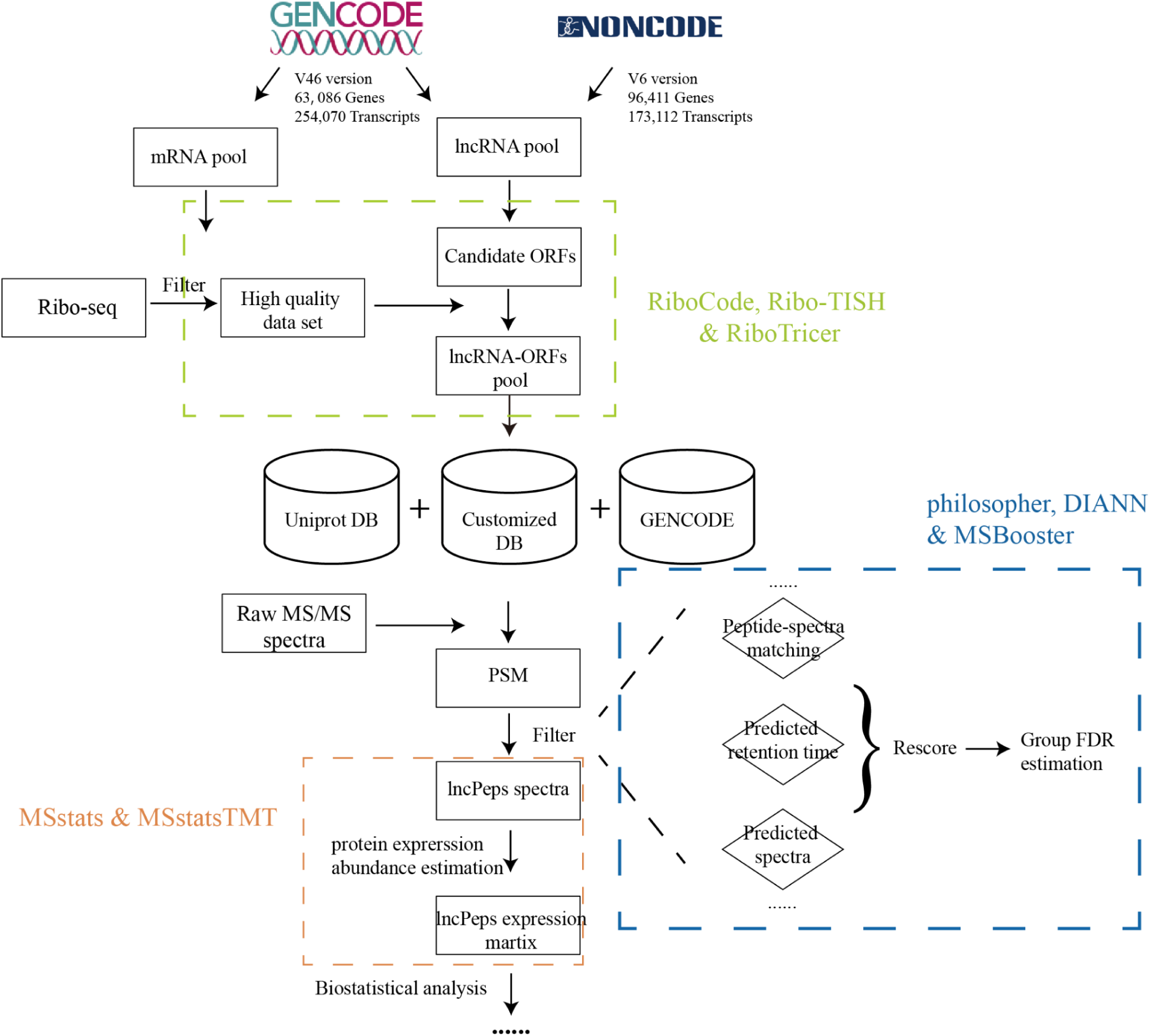
Supplementary figure for Figure 1. Workflow of lncPep discovery in this study. High-quality Ribo-seq datasets were curated and annotated using GENCODE and NONCODE references. Predicted lncRNA-ORFs were used to guide proteomic analysis, enabling the identification of high-confidence lncPeps from HCC patient biopsy samples.

**Figure S3:**
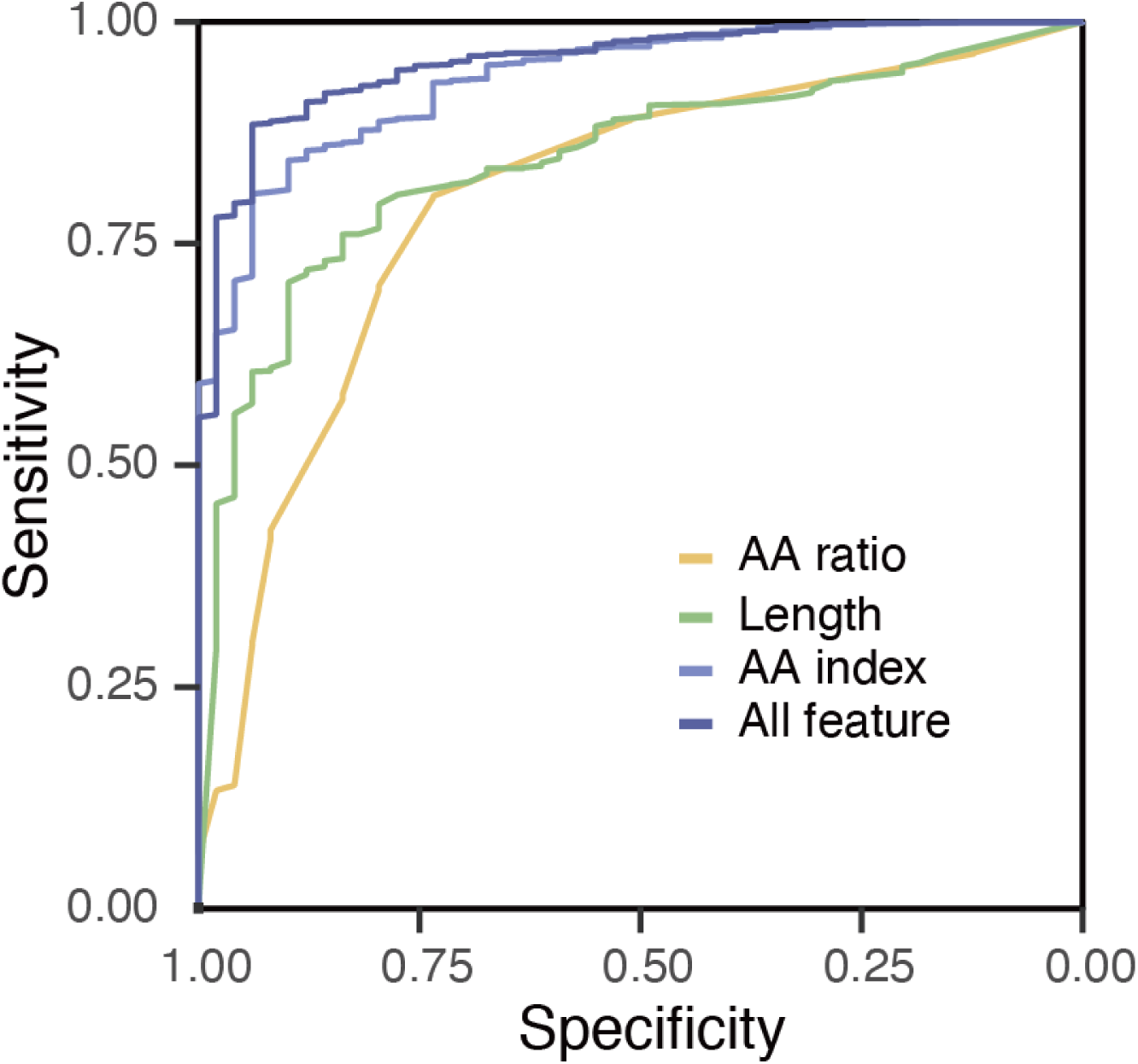
Supplementary figure for Figure 4. Prediction performance of machine learning models trained on amino acid composition, peptide length, AAindex features, or all features, evaluated on MS-peptides and Non-MS-peptides in the test set.

**Figure S4:**
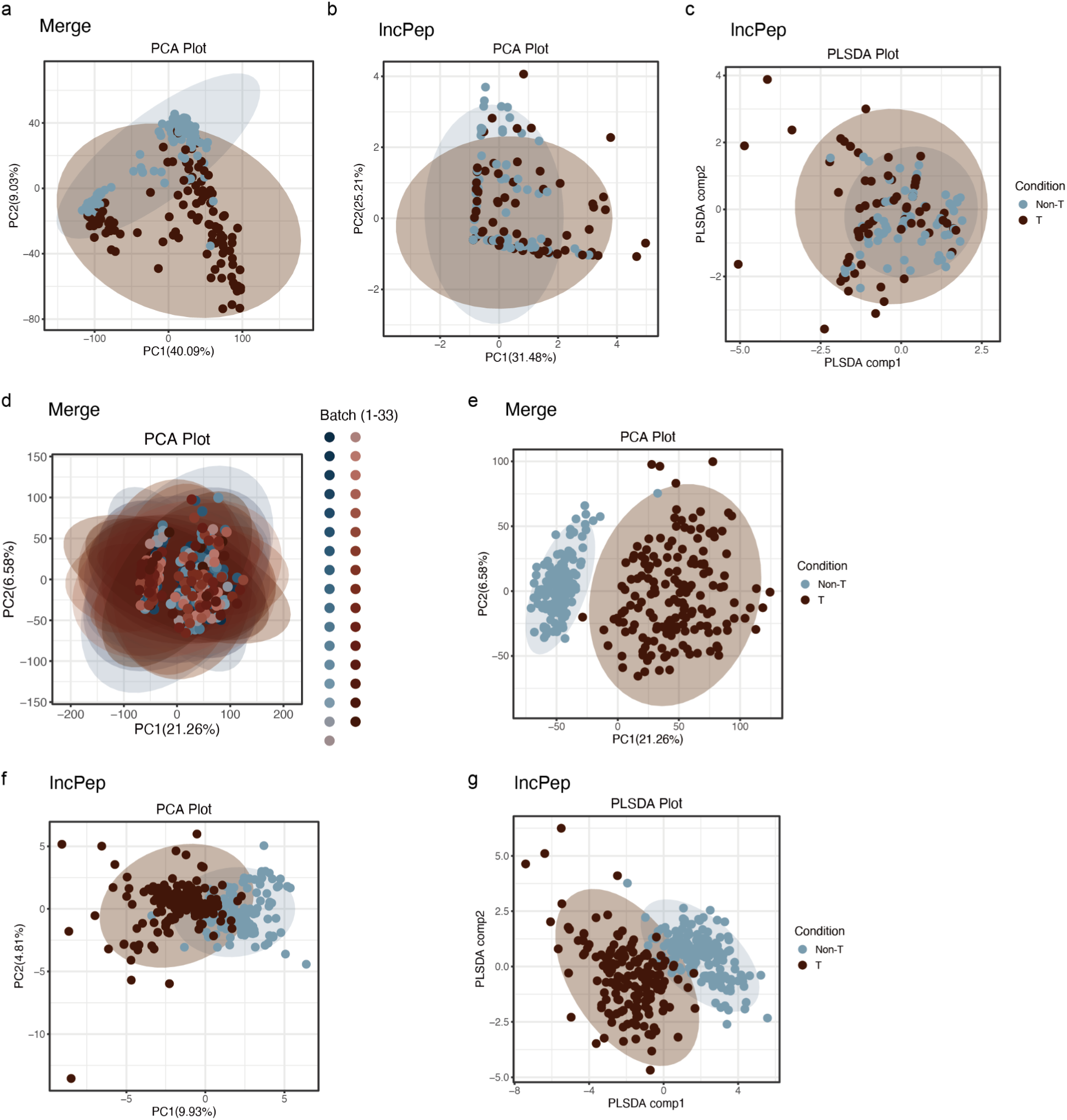
Supplementary figure for Figure 5. a–c. Principal component analysis (PCA) of the Merge (a) and lncPeps (b), and partial least squares discriminant analysis (PLS-DA) of lncPeps (c) in the Jiang et al. 2019 dataset. Non-T: Non-tumor and T: tumor tissues. d–g. Principal component analysis (PCA) and partial least squares discriminant analysis (PLS-DA) of the Gao et al. 2019 dataset. PCA of the Merge (d, e), PCA of lncPeps (f), and PLS-DA of lncPeps (g). Samples are labeled 01–33, corresponding to different TMT experimental batches.

**Figure S5:**
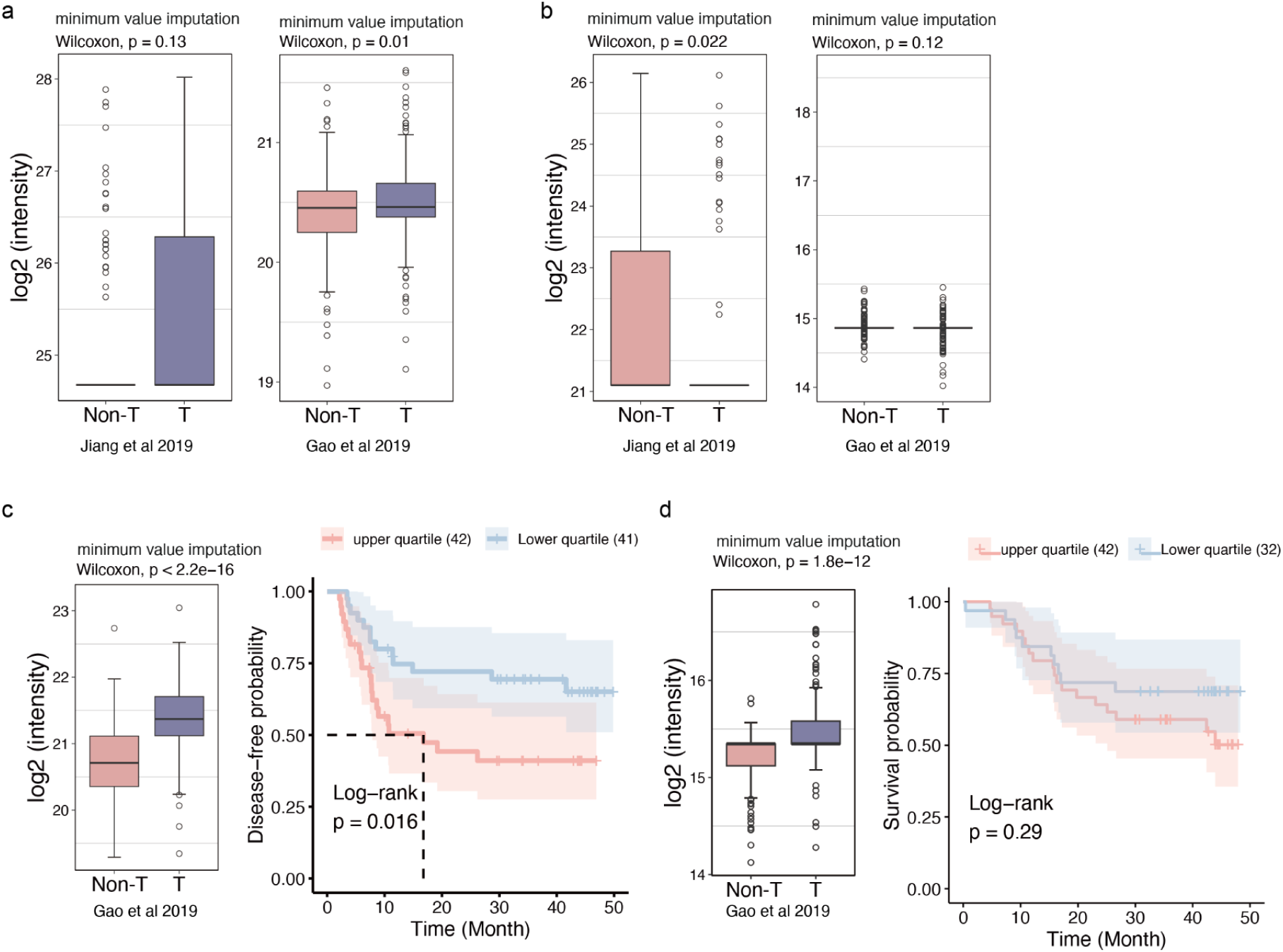
Supplementary figure for Figure 6. a–b. Box plots showing expression of *PPIAP79* lncORF1 (a) and *SEPTIN7P8* lncORF1 (b) in the Jiang et al. 2019 and Gao et al. 2019 datasets. Minimum value imputation was applied. Non-T: non-tumor, T: tumor tissues. c. Box plot showing expression of *POTEKP* lncORF2 in the Gao et al. 2019 dataset, and Kaplan–Meier (KM) analysis of survival probability for patients stratified by *POTEKP* lncORF2 expression (upper vs. lower quartile). Minimum value imputation was applied. d. Box plot showing expression of *HNRNPA1P36* lncORF1 in the Gao et al. 2019 dataset, and Kaplan–Meier (KM) analysis of survival probability for patients stratified by *HNRNPA1P36* lncORF1 expression (upper vs. lower quartile). Minimum value imputation was applied.

**Figure S6:**
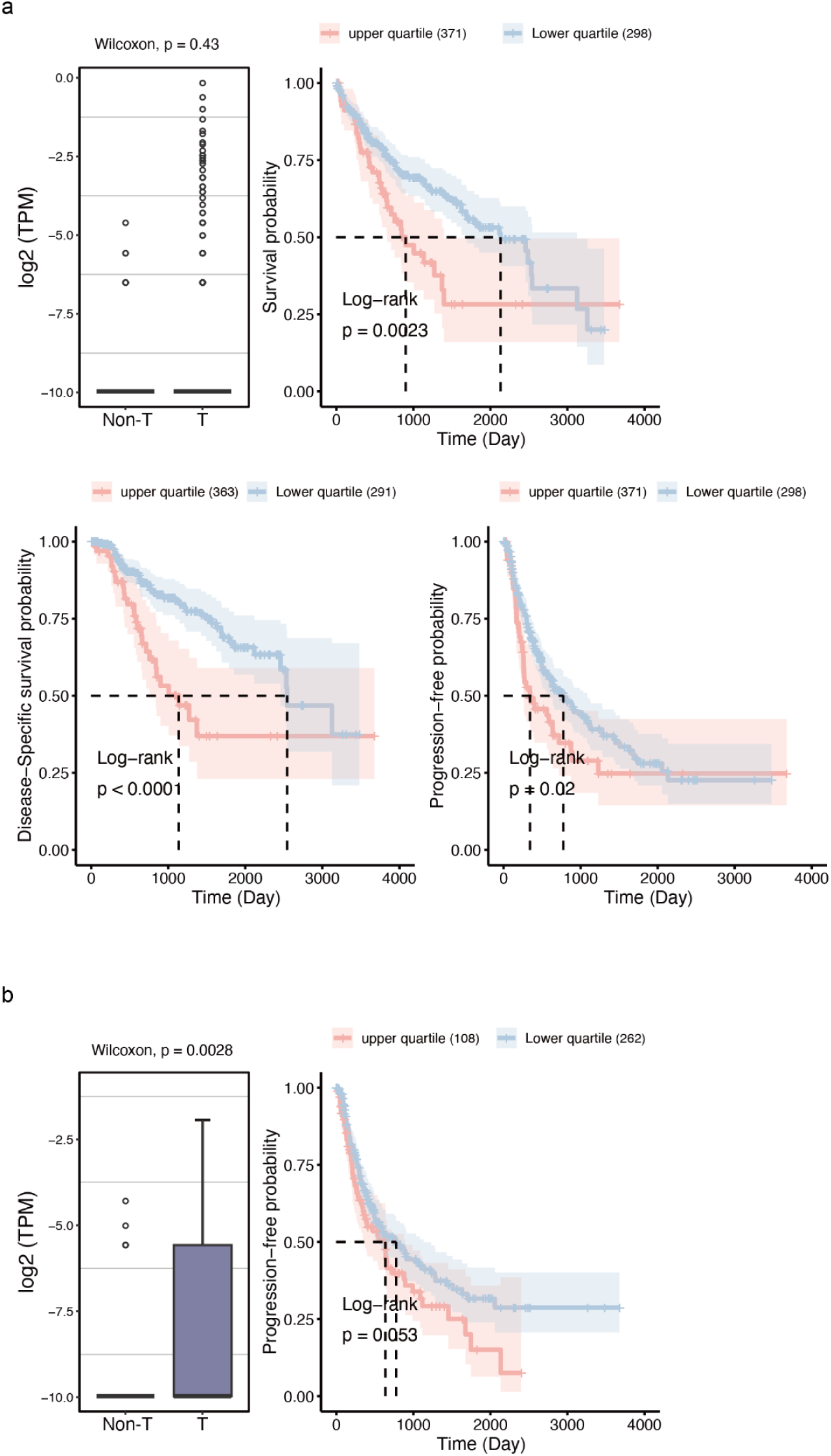
Supplementary figure for Figure 6. a. Box plot showing POTEKP RNA expression in non-tumor (Non-T) and tumor (T) samples from the TCGA dataset, and Kaplan–Meier (KM) analysis of overall survival (OS), disease-specific survival (DSS), and progression-free interval (PFI) for patients stratified by POTEKP expression (upper vs. lower quartile). b. Box plot showing HNRNPA1P36 RNA expression in the TCGA dataset, and KM analysis of progression-free interval (PFI) for patients stratified by HNRNPA1P36 expression (upper vs. lower quartile).

**Figure S7:**
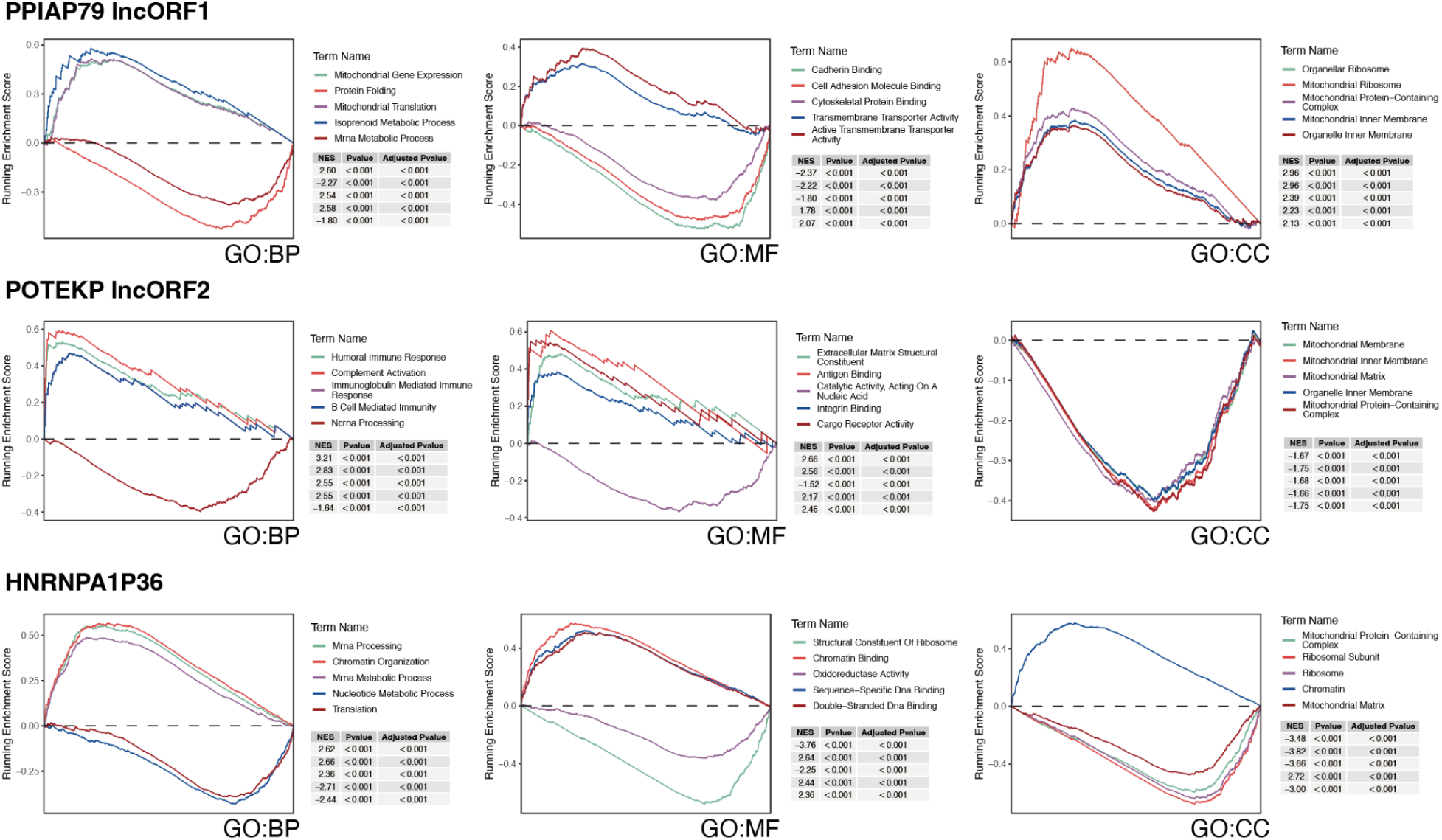
Supplementary figure for Figure 6. GSEA enrichment analysis of proteins associated with the expression of *PPIAP79* lncORF1, *POTEKP* lncORF2, and *HNRNPA1P36* lncORF1. Pearson correlation coefficients ranked proteins with lncPep expression, and enrichment results are shown for Gene Ontology (GO) categories: Biological Process (BP), Molecular Function (MF), and Cellular Component (CC).

